# Choosing Your Battles: Which Resistance Genes Warrant Global Action?

**DOI:** 10.1101/784322

**Authors:** An-Ni Zhang, Li-Guan Li, Xiaole Yin, Chengzhen L Dai, Mathieu Groussin, Mathilde Poyet, Edward Topp, Michael R Gillings, William P Hanage, James M Tiedje, Eric J Alm, Tong Zhang

## Abstract

The increasing accumulation of antibiotic resistance genes (ARGs) in pathogens poses a severe threat to the treatment of bacterial infections. However, not all ARGs do not pose the same threats to human health. Here, we present a framework to rank the risk of ARGs based on three factors: “anthropogenic enrichment”, “mobility”, and “host pathogenicity”. The framework is informed by all available bacterial genomes (55,000), plasmids (16,000), integrons (3,000), and 850 metagenomes covering diverse global eco-habitats. The framework prioritizes 3% of all known ARGs in Rank I (the most at risk of dissemination amongst pathogens) and 0.3% of ARGs in Rank II (high potential emergence of new resistance in pathogens). We further validated the framework using a list of 38 ARG families previously identified as high risk by the World Health Organization and published literature, and found that 36 of them were properly identified as top risk (Rank I) in our approach. Furthermore, we identified 43 unreported Rank I ARG families that should be prioritized for public health interventions. Within the same gene family, homologous genes pose different risks, host range, and ecological distributions, indicating the need for high resolution surveillance into their sequence variants. Finally, to help strategize the policy interventions, we studied the impact of industrialization on high risk ARGs in 1,120 human gut microbiome metagenomes of 36 diverse global populations. Our findings suggest that current policies on controlling the clinical antimicrobial consumptions could effectively control Rank I, while greater antibiotic stewardship in veterinary settings could help control Rank II. Overall, our framework offered a straightforward evaluation of the risk posed by ARGs, and prioritized a shortlist of current and emerging threats for global action to fight ARGs.

## Main

Antibiotic resistance genes (ARGs) are widespread pollutants^1–5^ (Figure S1) threatening human health. The World Health Organization (WHO) calls for global action to fight them^6^, but action cannot realistically be taken against thousands of known (and unknown) ARGs^7–9^. The risk of ARGs to human health varies considerably according to a number of factors including their genetic context. For example, intrinsic colistin resistance ARGs were found decades ago^10,11^ but their low potential for horizontal gene transfer (HGT) limited their spread. In contrast, the *mcr-1* colistin resistance gene has rapidly spread into eight pathogenic species (“host pathogenicity”)^12^ across 31 countries and into 1-20% animal and human gut microbiome samples^13–16^ (“anthropogenic enrichment”), largely driven by its capacity for HGT (“mobility”)^17^. These characteristics are typical of high risk ARGs, but current databases and analyses^7–9,18^ do not distinguish ARGs based on them. Mobilizing policymakers to implement interventions on ARGs will require substantial scientific capital, and prioritizing high risk ARGs will allow that capital to be effectively invested.

Therefore, there exists a need for a framework to categorize the risk of ARGs to human health. Previous attempts^19,20^ at defining such a framework based on critical factors have largely remained theoretical (such as mobility and sub-inhibitory antibiotic concentrations), and thus, difficult to implement. In this study, we developed an empirical approach that combines three factors (“anthropogenic enrichment”; “mobility”, and “host pathogenicity”) to prioritize our efforts and identify emerging threats.

## Results

ARGs enriched by anthropogenic activities, and correlated with antibiotics contamination should pose a higher risk than ARGs that are not enriched. Through investigating ARGs in 854 global metagenomes, we found that ARG composition clustered samples into three habitats along the primary axis of anthropogenicity (Figure 1a) from undisturbed natural habitats, to wastewater treatment plants (WWTPs), to feces. Through a literature review of 30 studies, we observed a 100-fold difference (p < 0.01 by kw-test) in the total concentration of antibiotics along the anthropogenicity axis (Figure 1b), posing a potential selection for the transmission and evolution of ARGs^21–25^ that increases bacterial fitness in human-related environments. These observations guided the framework that the anthropogenic impact (i.e., via antibiotics) primarily shaped the ARG composition, and should be assessed first to evaluate ARG risk. Yet the majority of ARGs^26 27^ were not impacted by anthropogenicity with no significant difference across habitats (p > 0.05 by kw-test, Figures 1b and S2a). This indicates that the genes we classify as ARGs probably have other primary functions in the environment^28^. Thus, we prioritize ARGs that are: enriched in anthropogenic environments (Figures 1b and S2b); are more likely to deal with clinically relevant antibiotics; and could be controlled by public health interventions that limit antibiotic usage.

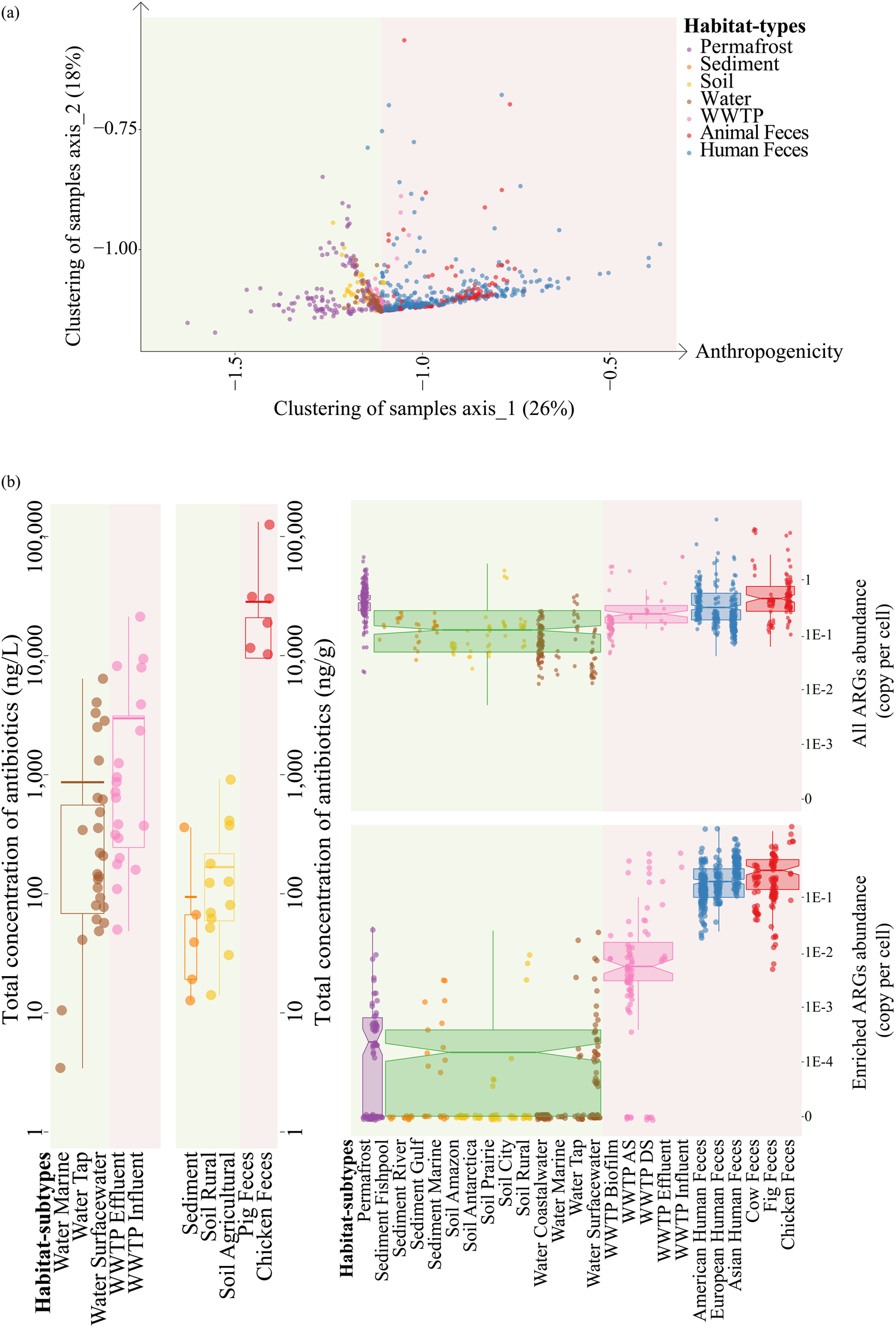

Our risk framework reflected the evolution and emergence of ARGs into human pathogens driven by antibiotics selection (Figure 2a). After assessing the anthropogenic enrichment, we prioritized mobile ARGs that are usually specialized resistance loci responding to antibiotics^21,29^ and are more capable of transferring between lineages^30–34^. We further prioritized ARGs carried by human pathogens, since this indicates a potential ineffective treatment of diseases by antibiotics. Based on these properties, we designed a risk framework that sequentially assessed three criteria (Figure 2b): Namely: being 100-fold more abundant in human dominated ecosystems than other ecosystems; being carried via mobile genetic elements (MGEs); and being resident in pathogens (see supplementary information and Figure S3 for analyses). We ranked all 4,050 ARGs in the Structured ARG Database^8^ (Table S2) and obtained relevant data by searching the ARGs in all available bacterial genomes and plasmids from NCBI, MGEs databases^35,36^, and 854 global metagenomes by the ARGs Online Searching Platform^37^. We found that 75% ARGs were not significantly responsive to anthropogenic impact (Rank IV) and were generally non-mobile (83%) and 5-10 fold more abundant in nature (Figure S4). Of all the anthropogenically enriched ARGs, 80% were non-mobile as Rank III ARGs. Rank III largely represented the intrinsic resistance evolved in pathogens as 80% of Rank III ARGs were hosted only by pathogens (50% by ESKAPE pathogens). We found that most mobile ARGs (132 of 145) were already carried by human pathogens as top risk Rank I ARGs. We consider Rank I as an immediate and current threat because they have several dangerous characteristics, such as notable host pathogenicity (93% in ESKAPE pathogens); wide host range (98% across several genera); and highly prevalent in impacted environments (14%) (Table S2). We also found 13 mobile ARGs that have not yet been carried by any pathogen (Rank II), which could pose a future threat through a potential contact and transfer into a pathogen.

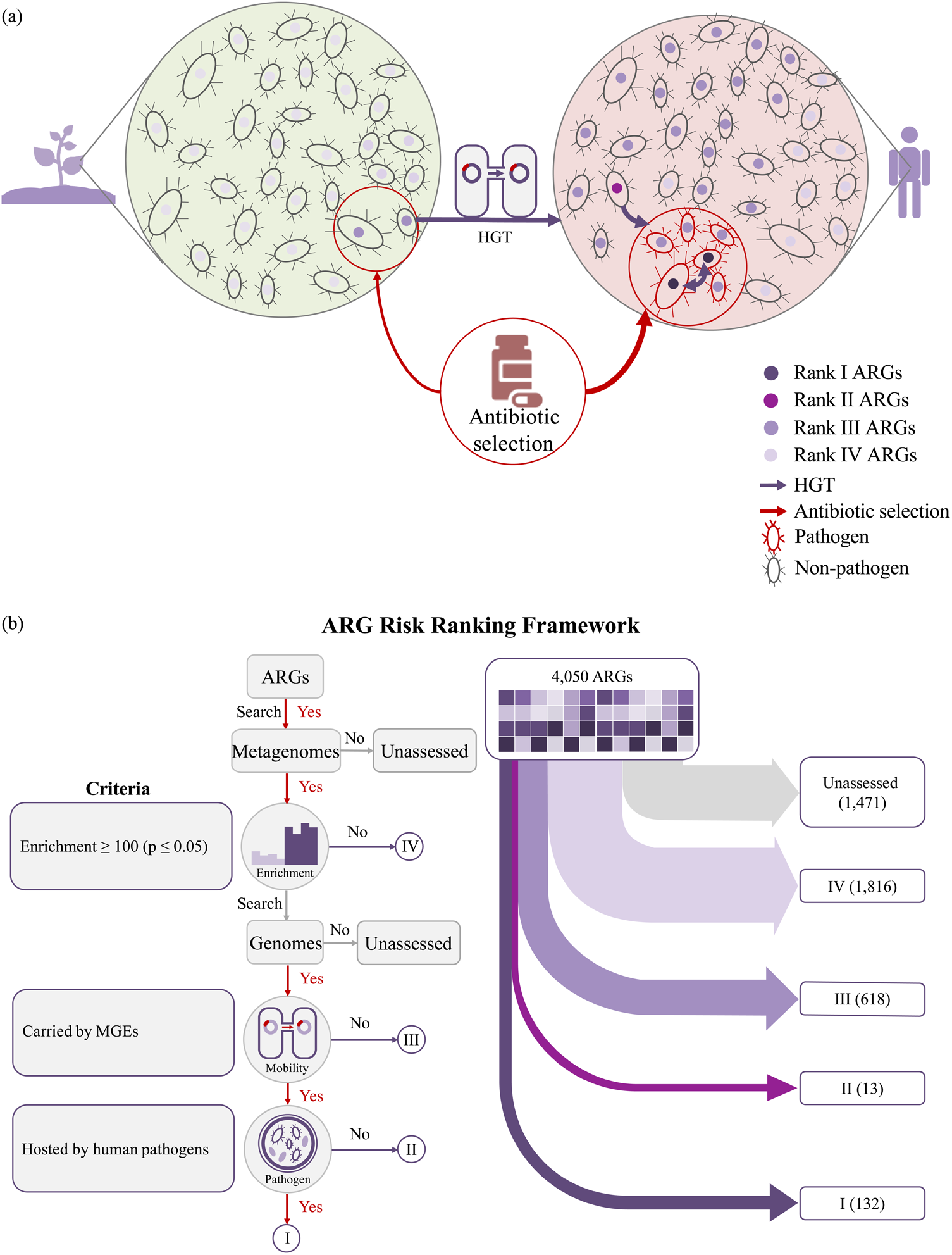

The ranking framework accurately identified 95% of all well-known high risk ARG families as Rank I ARGs and expanded the list by 43 unreported Rank I ARG families. We compared our results against a list of 38 ARG families that were reported to have high clinical concern (to cause treatment failure of healthcare associated infections and/or have been wide-spread phylogenetically and geographically) by the WHO and by literature review (highlighted in purple)^17,19,20,22,38–47^ (Figure 3). For example, we included extended-spectrum beta-lactamases (ESBL) ARGs (i.e., *TEM, SHV*, and *CTX-M*), that recently caused the death of fecal microbiota transplantation (FMT) recipients. Our framework properly identified 36 as Rank I and two as Rank IV. Both Rank IV ARGs (*sul1, vanA*) met the requirements of “mobility” and “pathogenicity”, but they were 5-20 fold more abundant in nature. More importantly, we automatically identified 43 Rank I ARG families that have not been reported as high risk in previous studies, but which exhibited the same features as frequently reported ARGs. Some of them have shown a strong clinical relevance (such as *IMP-4, OXA*^47–49^) and should also be prioritized for interventions. Rank I ARGs had a wide host range, in that 80% were shared across species and 60% across genera (same sequence variant), while 50% of Rank II-IV ARGs had a narrow host range within single species. We found homologs of one ARG family tended to pose different risks, different host range, and different ecological distributions. Usually only a few Rank I homologs covered a broad host range and others Rank I homologs shared a narrow subset (i.e., 14-16 genera compared to 1-3 genera for *tetM*). However, Rank II ARGs usually did not share the same host range with Rank I homologs (i.e., probiotic bacteria *Lactobacillus* for *tetM*). We also found Rank II ARGs without Rank I homologs and they were hosted by abundant gut commensals and/or close relatives to pathogens that could be a reservoir of new resistance for gut pathogens^50^, especially *aadA* and *vatE* with a host range across two orders. Besides, Rank I-II homologs did not share the same host strain (founder effect) and have distinct sequence variants in different habitats (Figure S5), while multiple Rank III-IV homologs within one genome were quite common. Thus, research and surveillance into ARGs should be conducted at the resolution of their sequence variants, not by just documenting their ARG families.

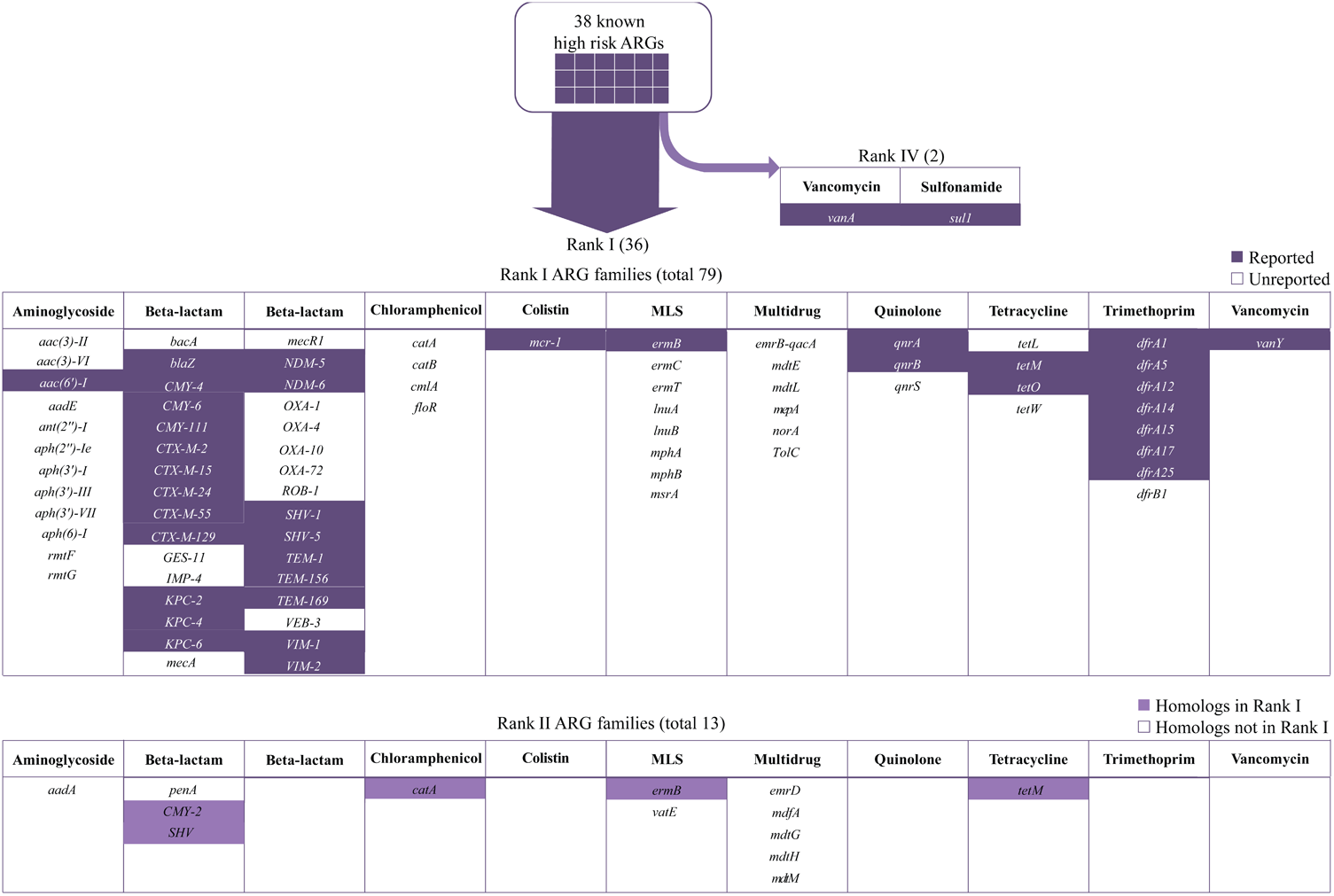

We observed that Rank I was strongly correlated with the potential exposure of clinical antibiotics, while Rank II was associated with industrialized lifestyles. We applied the ranking framework to 1,120 human gut microbiome metagenomes (36 global populations ref in preparation and 85 healthy FMT donors^51^) to understand how industrialization impacts the risk of antibiotic resistance. Industrialization dominated over all other factors that shaped the ARG composition across populations (Figure S6). Thus, we classified populations into five levels from non-industrialized rural lifestyles to industrialized urban lifestyles (details in Figure 4).

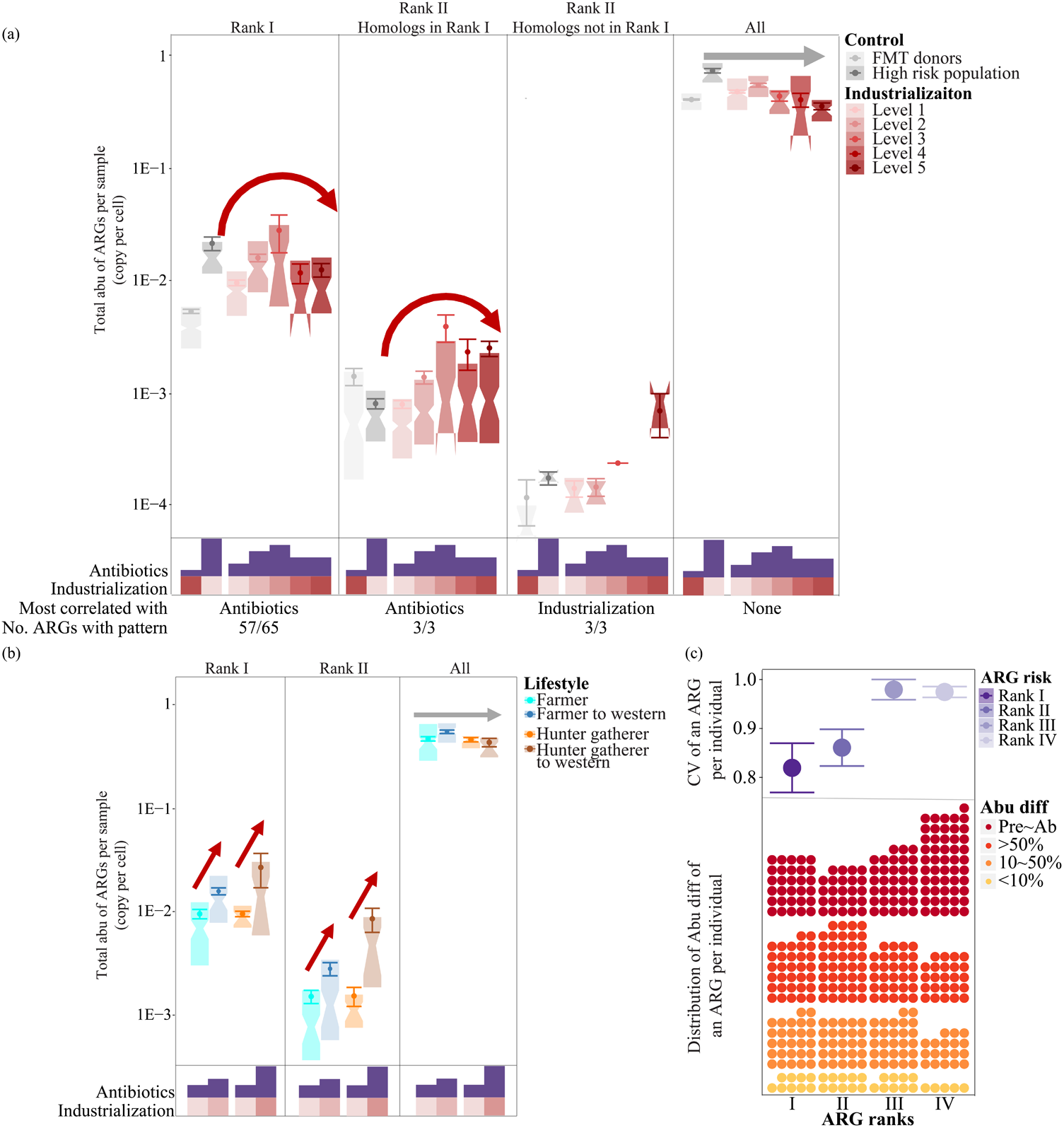

We roughly estimated the potential exposure to clinical antibiotics considering the access to antibiotics and policy interventions on antimicrobial consumptions^52–54^. Rank I was highly responsive to industrialization and revealed the same pattern with antibiotics exposure. In total of 57 of 65 Rank I ARGs presented this pattern individually (including *mcr-1* and all multidrug resistance), suggesting that this response was robust regardless of different host ranges and resistance mechanisms (Figure S8). Total Rank I was similarly abundant and diverse in non-industrialized rural populations (Level 1 with limited access with antibiotics) and highly industrialized populations (Levels 4-5 with policy interventions on clinical antimicrobial consumptions), and was significantly lower than middle industrialized populations (Levels 3-4 with access but limited interventions on clinical antimicrobial consumptions)^55^. Moreover, Rank I showed a 5-10 fold lower abundance (p < 0.01 by kw-test) in FMT donors who had no antibiotic consumptions from 6 months before sampling (Figure 4a). Meanwhile Rank I could maintain its low abundance with low variance in four FMT donors over 150-550 days of surveillance (Figures 4c and S8). These observations suggest that Rank I could be mainly associated with the potential exposure of clinical antibiotics. In addition, we detected 6 Rank II ARGs commonly shared across populations. Three Rank II ARGs with Rank I homologs (*catA, ermB, tetM*) were loosely correlated with clinical antibiotics. Since they were not carried by pathogens, they could be under a less selection than their Rank I homologs that were carried by pathogens. Another three novel Rank II ARGs (*aadA, vatE, mdtM*) were found highly abundant in the highest industrialized population (Level 5). Overall, Rank II was more abundant in industrialized populations (Levels 4-5) than high-risk population (pastoralist population that uses antibiotics in animal farms) and non-industrialized rural populations (Level 1), and most Rank II was less likely to get lost over time in FMT donors without antibiotic selection (Figure 4c). We also observed the transfer of lifestyles from farmers and hunter gatherers to industrialized lifestyles increased the Rank II by 3-10 fold (Figure 4b). Thus, Rank II seemed not to be correlated with clinical antibiotics but industrialized lifestyles. Instead, total ARGs displayed no significant difference across populations.

## Discussion

We designed a risk-ranking framework to prioritize high risk ARGs through “anthropogenic enrichment”, “mobility”, and “host pathogenicity”. Our framework successfully identified 36 of 38 high risk ARG that have been reported (including *mcr-1*) and expanded the list by 43 unreported emerging Rank I ARGs. Management strategies should be evaluated and implemented on the basis of their ability to control these high risk ARGs. Moreover, the surveillance of high risk ARGs should target specific sequence variants since homologs (homologous ARGs) in the same ARG family pose different risks and different phylogenetic and ecological preference. The representativeness of available public datasets is the primary limitation of this study. With new sequencing data and clinical data, this ranking framework is expected to supervise the dynamic status of ARG risk and to identify future threats. Finally, to help strategize public health interventions, we studied the impact of industrialization and antibiotics exposure on high risk ARGs in global human gut microbiome. We found that current policy controlling the clinical antimicrobial consumptions could effectively control Rank I (current threat)^56,57^ but not Rank II (future threat). We observed that antibiotics exposure could explain the pattern of Rank I, especially its relatively low diversity and abundance in industrialized populations with strict policy interventions on clinical antimicrobial consumptions^54^. Besides, Rank I was maintained at a significantly low abundance in FMT donors with no antibiotic consumptions. These observations suggested that Rank I could strongly and rapidly respond to clinical antibiotics^56^ and was effectively controlled by current policy in developed countries^54^ (such as therapy guidelines, antibiotic formularies, antibiotic stewardship programmes, and public health interventions^58^). However, we found that Rank II was not directly correlated with the clinical antibiotics and was not observed to be effectively controlled in industrialized countries. The fact that novel Rank II ARGs were usually carried by abundant gut commensals and/or close relatives to gut pathogens, indicated their high potential and consequences to emerge into gut pathogens. We should take actions with new intervention strategies, such as controlling the usage of veterinary antibiotics as growth promoters^59^ (i.e, *vatE*)^60,61^.

## Methods

Details of methods, data and scripts are all available in supplementary information. Briefly, the ARGs Online Searching Platform^37^ provided the presence and abundance of ARGs in 54,718 all available NCBI bacterial genomes (≥ 50% completeness and < 10% contamination) and 854 global metagenomes of Illumina shotgun sequencing. We further searched ARGs in all available NCBI 15,738 plasmids and other mobile genetic element (MGE) databases^35,36^ by usearch v11.0, diamond 0.9.24 and blast 2.5.0+^62–64^. The search cutoff was set for genomes and MGEs as e-value 1e-5, 90% aa similarity over 80% aa hit length; and for metagenomes as e-value 1e-7, 80% aa similarity over 75% aa hit length^7,8,37,65^. The abundance of ARGs was normalized into copy of genes per bacterial cell (ARGs-OAP^7,8^) using Equation 1. The total bacterial cell number of one metagenomic sample was inferred by counting the average copy number of bacterial essential single copy genes.

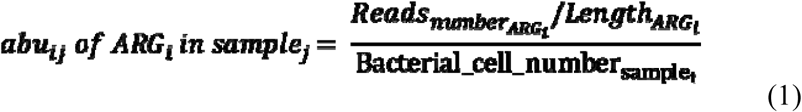

We developed a bioinformatics tool (arg_ranker) to assess the risk of ARGs in metagenomes and genomes (https://github.com/caozhichongchong/arg_ranker). The arg_ranker classifies all ARGs in one sample into Rank I-IV and quantifies the risk contributed by each Rank.

We used 460 human gut microbiome metagenomes from 460 donors covering 36 different populations of 9 lifestyles (ref in preparation) and 560 metagenomes from 84 FMT donors^51^. The FMT metagenomes consisted of 400 metagenomes of four donors with intensive sampling (231 samples over 536 days, 83 samples over 375 days, 70 samples over 201 days, and 90 samples over 144 days) and 160 metagenomes for 80 donors with sparse sampling (2 samples per individual over 2 to 460 days). We classified the populations into five industrialization level and we roughly estimated the potential exposure to antibiotics based on the access to antibiotics and public health interventions on clinical antimicrobial consumptions of each population^52–54^. FMT donors are healthy individuals with no antibiotic consumptions from six months before sampling, which represents the least potential exposure to antibiotics. The high-risk population is a pastoralist population that uses antibiotics in animal farms, which represents the highest potential exposure to antibiotics. Level 1: non-industrialized rural populations with limited access to antibiotics (fisherman, hunter gatherer to farmer, farmer, and hunter gatherer); Level 2: non-industrialized urban populations (farmer to western) with access to antibiotics and little policy interventions^54^; Level 3: industrialized rural population with access to antibiotics and little policy interventions^54^ (hunter gatherer to western); Level 4: industrialized rural populations (western) with access to antibiotics and policy interventions; Level 5: industrialized urban populations (western) with access to antibiotics and policy interventions. The relative abundance difference (abu_diff_) of an ARG between 2 time-points in one individual was calculated using Equation 2.

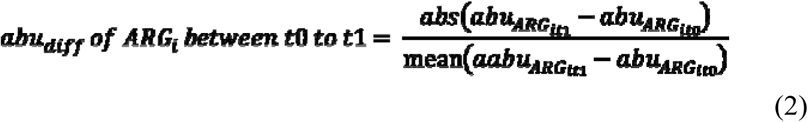

## Acknowledgements

Dr. A.N. Zhang, Dr. L.G Li, and X. Yin acknowledge the University of Hong Kong for the graduate and postgraduate studentships. Dr. A.N. Zhang and C.D. acknowledge the MIT for for the graduate and postgraduate studentships. The authors would like to thank the Hong Kong General Research Fund, Broad Institute (Broad Next 10 grant 4000017), and Center for Microbiome Informatics and Therapeutics at MIT for the financial support.

## Author contributions

A.N.Z. developed the pipeline, analyzed the data, and wrote the manuscript. L.L. contributed suggestions in data analysis and manuscript preparation. C.D. and X.Y. contributed suggestions on data analysis and revised the manuscript. M.G. and M.P. processed and provided gut microbiome metagenomic data, contributed suggestions on data analysis, and revised the manuscript. W.P.H., J.M.T., and E.T. provided valuable encouragement, contributed valuable advice on the pipeline development, and revised this manuscript. E.J.A. and T.Z. guided the pipeline development, data analysis, and revised this manuscript.

## References

1. D’Costa, V. M. et al. Antibiotic resistance is ancient. Nature 477, 457 (2011).

2. Hu, Y. et al. Metagenome-wide analysis of antibiotic resistance genes in a large cohort of human gut microbiota. Nat Commun 4, 2151 (2013).

3. Czekalski, N., Díez, E. G. & Bürgmann, H. Wastewater as a point source of antibiotic-resistance genes in the sediment of a freshwater lake. ISME J 8, 1381 (2014).

4. Tang, J. et al. Metagenomic analysis of bacterial community composition and antibiotic resistance genes in a wastewater treatment plant and its receiving surface water. Ecotoxicol Env. Saf 132, 260–269 (2016).

5. Lau, C. H.-F., van Engelen, K., Gordon, S., Renaud, J. & Topp, E. Novel antibiotic resistance determinants from agricultural soil exposed to antibiotics widely used in human medicine and animal farming. Appl Env. Microb. AEM. 00989-17 (2017).

6. Antimicrobial resistance: global report on surveillance. (World Health Organization, 2014).

7. Yin, X. et al. ARGs-OAP v2. 0 with an Expanded SARG Database and Hidden Markov Models for Enhancement Characterization and Quantification of Antibiotic Resistance Genes in Environmental Metagenomes. Bioinformatics 1, 8 (2018).

8. Yang, Y. et al. ARGs-OAP: online analysis pipeline for antibiotic resistance genes detection from metagenomic data using an integrated structured ARG-database. Bioinformatics (2016). doi:10.1093/bioinformatics/btw136

9. Jia, B. et al. CARD 2017: expansion and model-centric curation of the comprehensive antibiotic resistance database. Nucleic Acids Res gkw1004 (2016).

10. Olaitan, A. O., Morand, S. & Rolain, J.-M. Mechanisms of polymyxin resistance: acquired and intrinsic resistance in bacteria. Front. Microbiol. 5, 643 (2014).

11. Nordmann, P., Jayol, A. & Poirel, L. Rapid detection of polymyxin resistance in Enterobacteriaceae. Emerg. Infect. Dis. 22, 1038 (2016).

12. Wang, R. et al. The global distribution and spread of the mobilized colistin resistance gene mcr-1. Nat. Commun. 9, 1179 (2018).

13. Fernandes, M. R. et al. Silent dissemination of colistin-resistant Escherichia coli in South America could contribute to the global spread of the mcr-1 gene. Eurosurveillance 21, (2016).

14. Skov, R. L. & Monnet, D. L. Plasmid-mediated colistin resistance (mcr-1 gene): three months later, the story unfolds. Eurosurveillance 21, 30155 (2016).

15. McGann, P. et al. Escherichia coli harboring mcr-1 and blaCTX-M on a novel IncF plasmid: first report of mcr-1 in the United States. Antimicrob. Agents Chemother. 60, 4420–4421 (2016).

16. Rapoport, M. et al. First description of mcr-1-mediated colistin resistance in human infections caused by Escherichia coli in Latin America. Antimicrob. Agents Chemother. 60, 4412–4413 (2016).

17. Liu, Y.-Y. et al. Emergence of plasmid-mediated colistin resistance mechanism MCR-1 in animals and human beings in China: a microbiological and molecular biological study. Lancet Infect. Dis. 16, 161–168 (2016).

18. Liu, B. & Pop, M. ARDB—antibiotic resistance genes database. Nucleic Acids Res 37, D443–D447 (2008).

19. Martinez, J. L., Coque, T. M. & Baquero, F. What is a resistance gene? Ranking risk in resistomes. Nat Rev Microbiol 13, 116–23 (2015).

20. Berendonk, T. U. et al. Tackling antibiotic resistance: the environmental framework. Nat Rev Microbiol 13, 310–7 (2015).

21. Ji, X. et al. Antibiotic resistance gene abundances associated with antibiotics and heavy metals in animal manures and agricultural soils adjacent to feedlots in Shanghai; China. J Hazard Mater 235, 178–185 (2012).

22. Berglund, B. Environmental dissemination of antibiotic resistance genes and correlation to anthropogenic contamination with antibiotics. Infect. Ecol. Epidemiol. 5, 28564 (2015).

23. Gao, P., Munir, M. & Xagoraraki, I. Correlation of tetracycline and sulfonamide antibiotics with corresponding resistance genes and resistant bacteria in a conventional municipal wastewater treatment plant. Sci. Total Environ. 421, 173–183 (2012).

24. Pruden, A., Arabi, M. & Storteboom, H. N. Correlation between upstream human activities and riverine antibiotic resistance genes. Env. Sci Technol 46, 11541–11549 (2012).

25. Xu, J. et al. Occurrence of antibiotics and antibiotic resistance genes in a sewage treatment plant and its effluent-receiving river. Chemosphere 119, 1379–1385 (2015).

26. Webber, M. & Piddock, L. The importance of efflux pumps in bacterial antibiotic resistance. J. Antimicrob. Chemother. 51, 9–11 (2003).

27. Blair, J. M., Webber, M. A., Baylay, A. J., Ogbolu, D. O. & Piddock, L. J. Molecular mechanisms of antibiotic resistance. Nat. Rev. Microbiol. 13, 42 (2015).

28. Breidenstein, E. B., de la Fuente-Núñez, C. & Hancock, R. E. Pseudomonas aeruginosa: all roads lead to resistance. Trends Microbiol 19, 419–426 (2011).

29. Vaz-Moreira, I., Nunes, O. C. & Manaia, C. M. Bacterial diversity and antibiotic resistance in water habitats: searching the links with the human microbiome. FEMS Microbiol Rev 38, 761–778 (2014).

30. Nathwani, D., Raman, G., Sulham, K., Gavaghan, M. & Menon, V. Clinical and economic consequences of hospital-acquired resistant and multidrug-resistant Pseudomonas aeruginosa infections: a systematic review and meta-analysis. Antimicrob. Resist. Infect. Control 3, 32 (2014).

31. Alonso, A., Sanchez, P. & Martinez, J. L. Environmental selection of antibiotic resistance genes. Env. Microbiol 3, 1–9 (2001).

32. Baquero, F., Alvarez-Ortega, C. & Martinez, J. Ecology and evolution of antibiotic resistance. Env. Microbiol Rep 1, 469–476 (2009).

33. Martinez, J. L. The role of natural environments in the evolution of resistance traits in pathogenic bacteria. Proc R Soc Lond Ser B Biol Sci 276, 2521–2530 (2009).

34. Martínez, J. L. Natural antibiotic resistance and contamination by antibiotic resistance determinants: the two ages in the evolution of resistance to antimicrobials. Front Microbiol 3, 1 (2012).

35. Zhang, A. N. et al. Conserved phylogenetic distribution and limited antibiotic resistance of class 1 integrons revealed by assessing the bacterial genome and plasmid collection. Microbiome 6, 130 (2018).

36. Jiang, X., Hall, A. B., Xavier, R. J. & Alm, E. J. Comprehensive analysis of mobile genetic elements in the gut microbiome reveals a phylum-level niche-adaptive gene pool. bioRxiv 214213 (2017).

37. Zhang, A. N., Hou, C.-J., Li, L.-G. & Zhang, T. ARGs-OSP: online searching platform for antibiotic resistance genes distribution in metagenomic database and bacterial whole genome database. bioRxiv 337675 (2018).

38. Le, T.-H. et al. Occurrences and characterization of antibiotic-resistant bacteria and genetic determinants of hospital wastewater in a tropical country. Antimicrob. Agents Chemother. 60, 7449–7456 (2016).

39. French, G., Shannon, K. & Simmons, N. Hospital outbreak of Klebsiella pneumoniae resistant to broad-spectrum cephalosporins and beta-lactam-beta-lactamase inhibitor combinations by hyperproduction of SHV-5 beta-lactamase. J. Clin. Microbiol. 34, 358–363 (1996).

40. Feizabadi, M. M. et al. Distribution of bla TEM, bla SHV, bla CTX-M genes among clinical isolates of Klebsiella pneumoniae at Labbafinejad Hospital, Tehran, Iran. Microb. Drug Resist. 16, 49–53 (2010).

41. Coque, T. M., Oliver, A., Pérez-Díaz, J. C., Baquero, F. & Cantón, R. Genes encoding TEM-4, SHV-2, and CTX-M-10 extended-spectrum β-lactamases are carried by multiple Klebsiella pneumoniae clones in a single hospital (Madrid, 1989 to 2000). Antimicrob. Agents Chemother. 46, 500–510 (2002).

42. Yong, D. et al. Characterization of a new metallo-β-lactamase gene, blaNDM-1, and a novel erythromycin esterase gene carried on a unique genetic structure in Klebsiella pneumoniae sequence type 14 from India. Antimicrob Agents Chemother 53, 5046–5054 (2009).

43. Betteridge, T., Partridge, S. R., Iredell, J. R. & Stokes, H. W. Genetic context and structural diversity of class 1 integrons from human commensal bacteria in a hospital intensive care unit. Antimicrob Agents Chemother 55, 3939–43 (2011).

44. Lee, W. G., Jernigan, J. A., Rasheed, J. K., Anderson, G. J. & Tenover, F. C. Possible Horizontal Transfer of thevanB2 Gene among Genetically Diverse Strains of Vancomycin-Resistant Enterococcus faecium in a Korean Hospital. J. Clin. Microbiol. 39, 1165–1168 (2001).

45. Del Campo, R. et al. Detection of a single van A-containing Enterococcus faecalis clone in hospitals in different regions in Spain. J. Antimicrob. Chemother. 48, 746–747 (2001).

46. Valdezate, S. et al. Large clonal outbreak of multidrug-resistant CC17 ST17 Enterococcus faecium containing Tn 5382 in a Spanish hospital. J. Antimicrob. Chemother. 63, 17–20 (2008).

47. Karthikeyan, K., Thirunarayan, M. & Krishnan, P. Coexistence of bla OXA-23 with bla NDM-1 and armA in clinical isolates of Acinetobacter baumannii from India. J. Antimicrob. Chemother. 65, 2253–2254 (2010).

48. Peleg, A. Y., Franklin, C., Bell, J. M. & Spelman, D. W. Dissemination of the metallo-β-lactamase gene bla IMP-4 among gram-negative pathogens in a clinical setting in Australia. Clin. Infect. Dis. 41, 1549–1556 (2005).

49. Peleg, A. Y., Franklin, C., Bell, J. & Spelman, D. W. Emergence of IMP-4 metallo-β-lactamase in a clinical isolate from Australia. J. Antimicrob. Chemother. 54, 699–700 (2004).

50. Gueimonde, M., Sánchez, B., de los Reyes-Gavilán, C. G. & Margolles, A. Antibiotic resistance in probiotic bacteria. Front. Microbiol. 4, 202 (2013).

51. Poyet, M. et al. A library of human gut bacterial isolates paired with longitudinal multiomics data enables mechanistic microbiome research. Nat. Med. 1–11 (2019).

52. WHO Report on Surveillance of Antibiotic Consumption. (World Health Organization, 2018).

53. Geographical distribution of the consumption of Antibacterials for systemic use (ATC group J01) in the community (primary care sector) in Europe, reporting year 2017. (European Centre for Disease Prevention and Control, 2017).

54. Cox, J. et al. Antibiotic stewardship in low-and middle-income countries: the same but different? Clin. Microbiol. Infect. 23, 812–818 (2017).

55. Hendriksen, R. S. et al. Global monitoring of antimicrobial resistance based on metagenomics analyses of urban sewage. Nat. Commun. 10, 1124 (2019).

56. Dancer, S. et al. Approaching zero: temporal effects of a restrictive antibiotic policy on hospital-acquired Clostridium difficile, extended-spectrum β-lactamase-producing coliforms and meticillin-resistant Staphylococcus aureus. Int. J. Antimicrob. Agents 41, 137–142 (2013).

57. Cook, P. P. & Gooch, M. Long-term effects of an antimicrobial stewardship programme at a tertiary-care teaching hospital. Int. J. Antimicrob. Agents 45, 262–267 (2015).

58. Tacconelli, E. et al. Surveillance for control of antimicrobial resistance. Lancet Infect. Dis. 18, e99–e106 (2018).

59. Van Boeckel, T. P. et al. Global trends in antimicrobial resistance in animals in low- and middle-income countries. Science 365, (2019).

60. Wegener, H. C. Antibiotics in animal feed and their role in resistance development. Curr. Opin. Microbiol. 6, 439–445 (2003).

61. Werner, G. et al. Quinupristin/dalfopristin-resistant enterococci of the satA (vatD) and satG (vatE) genotypes from different ecological origins in Germany. Microb. Drug Resist. 6, 37–47 (2000).

62. Camacho, C. et al. BLAST+: architecture and applications. Bioinformatics 10, 421 (2009).

63. Edgar, R. C. Search and clustering orders of magnitude faster than BLAST. Bioinformatics 26, 2460–2461 (2010).

64. Buchfink, B., Xie, C. & Huson, D. H. Fast and sensitive protein alignment using DIAMOND. Nat Methods 12, 59 (2015).

65. Li, L.-G., Xia, Y. & Zhang, T. Co-occurrence of antibiotic and metal resistance genes revealed in complete genome collection. ISME J 11, 651–662 (2017).

