## Supplementary figures and images for "Choosing Your Battles: Which Resistance Genes Warrant Global Action?"

### Supplemental Figure 1

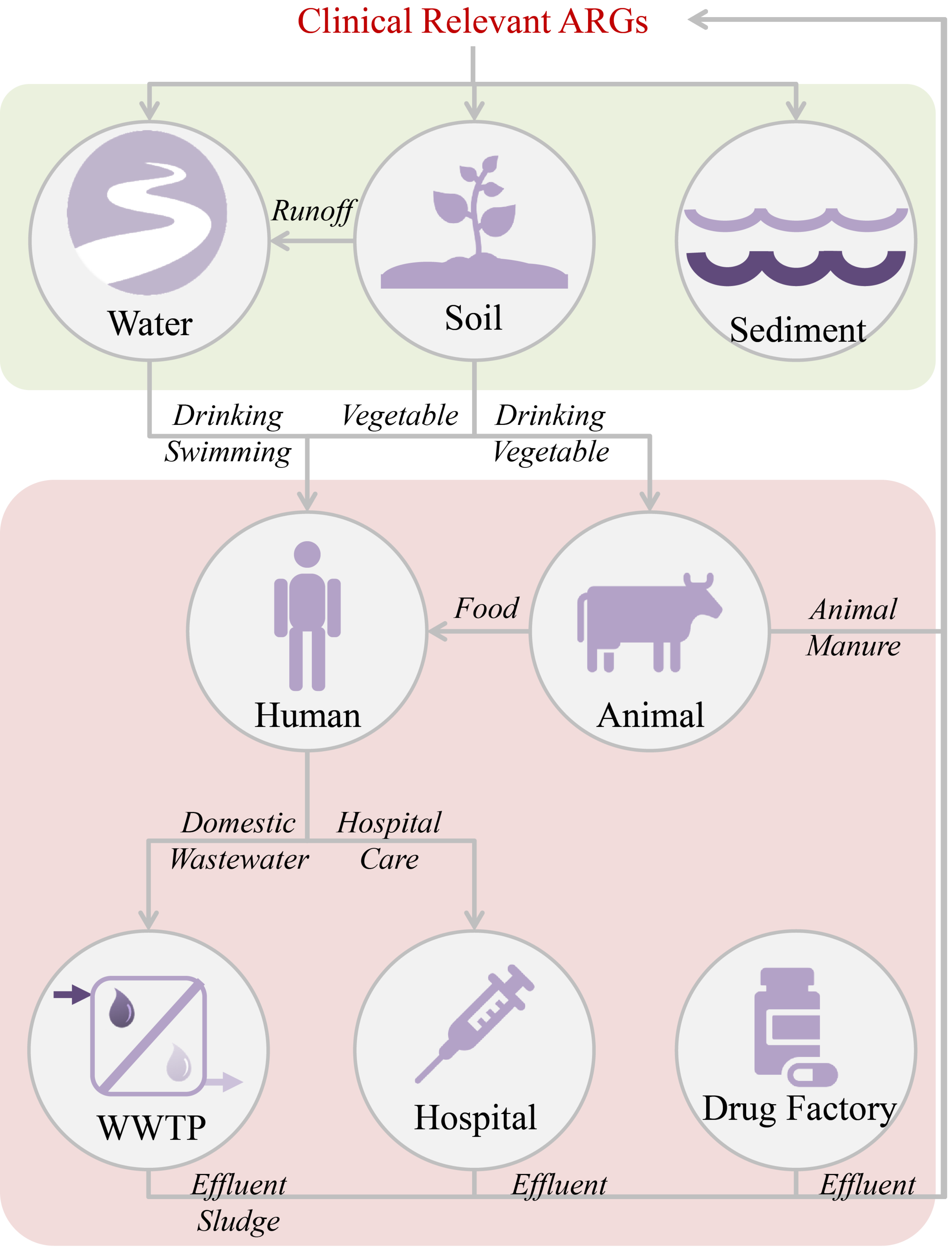

### Supplemental Figure 2

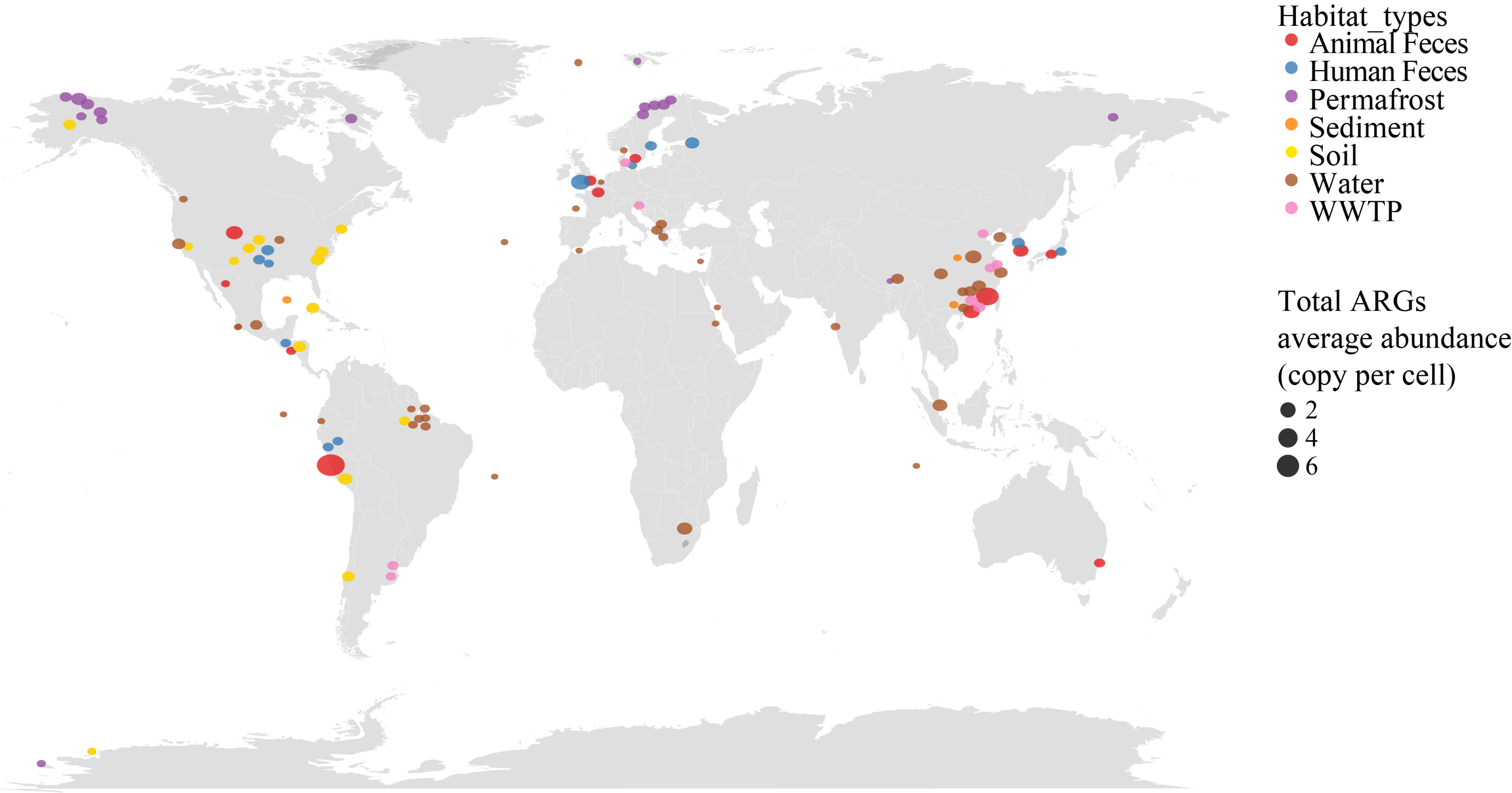

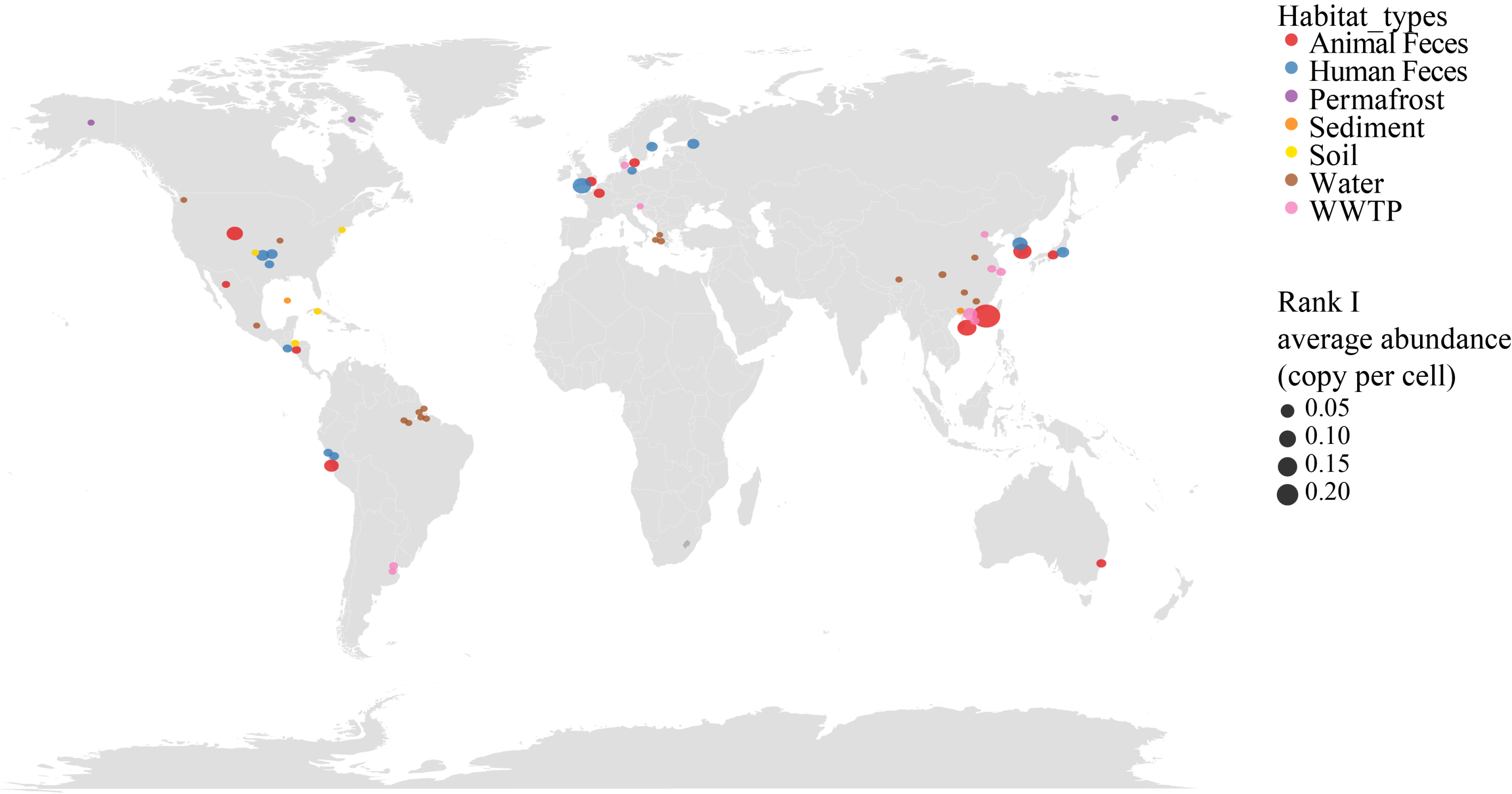

### Supplemental Figure 3

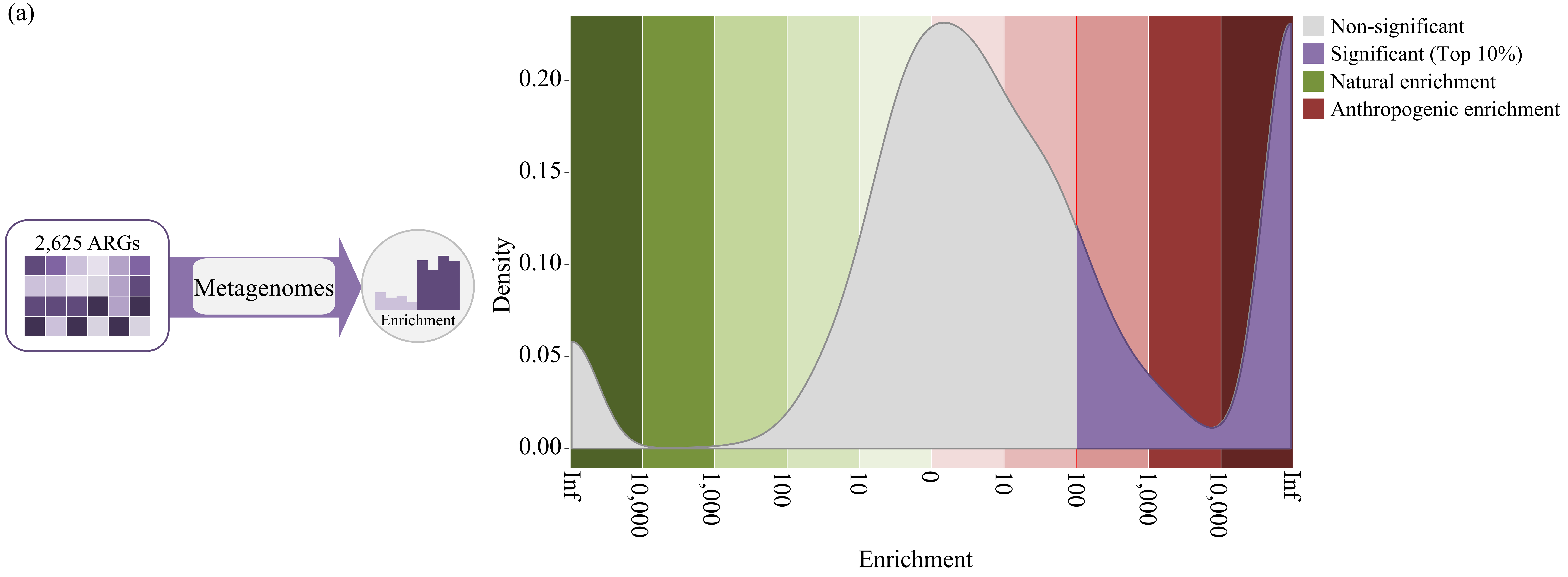

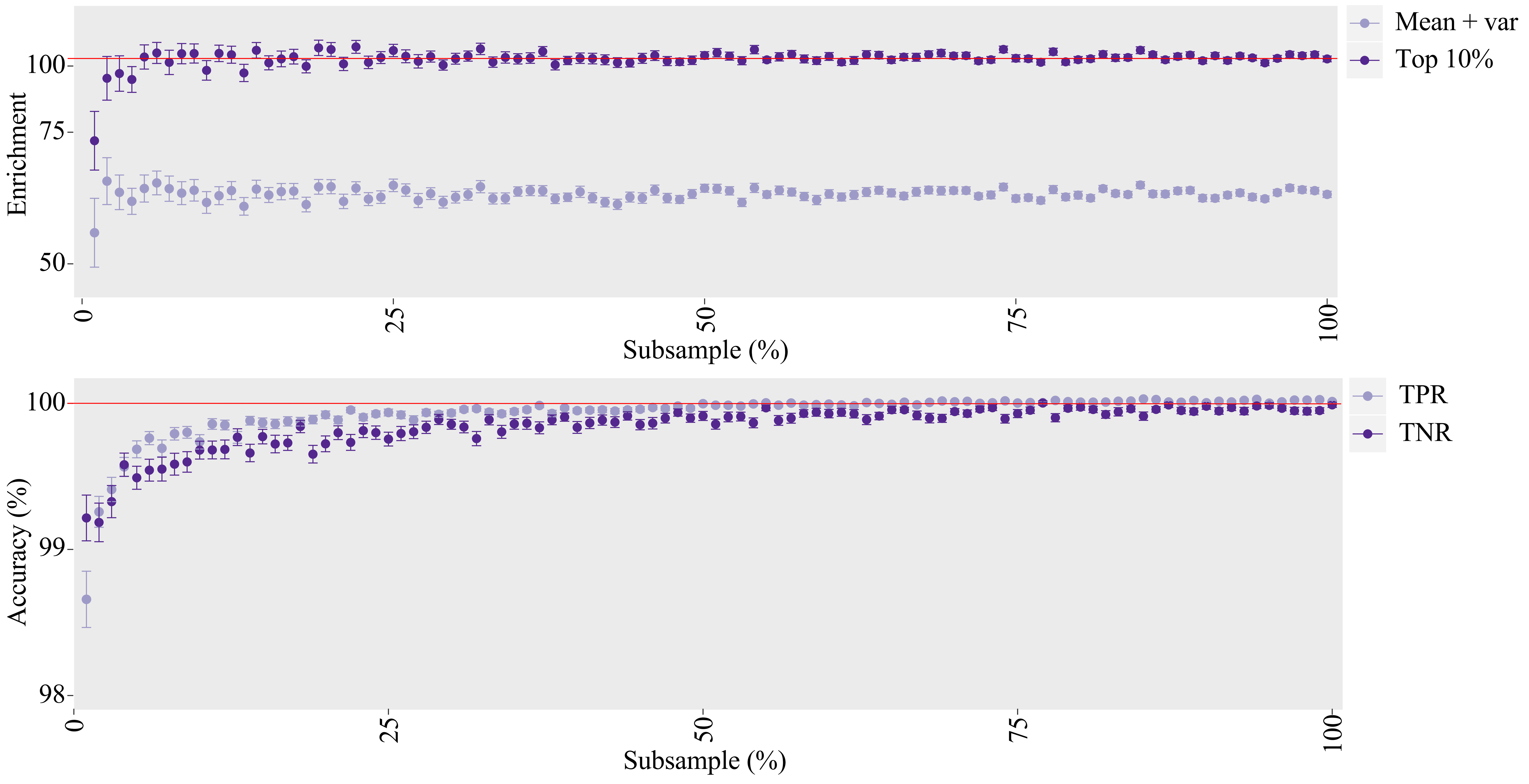

### Supplemental Figure 4

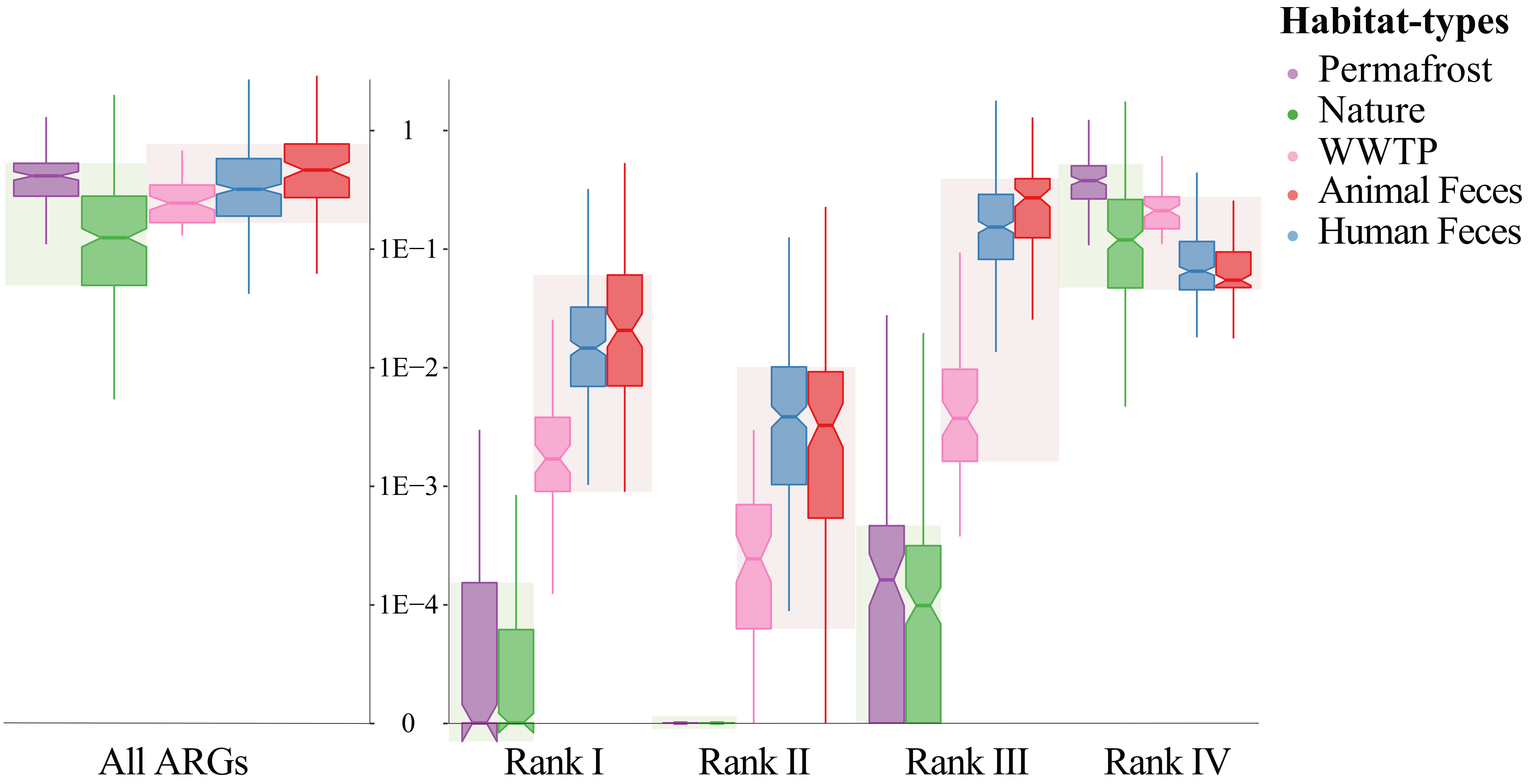

### Supplemental Figure 5

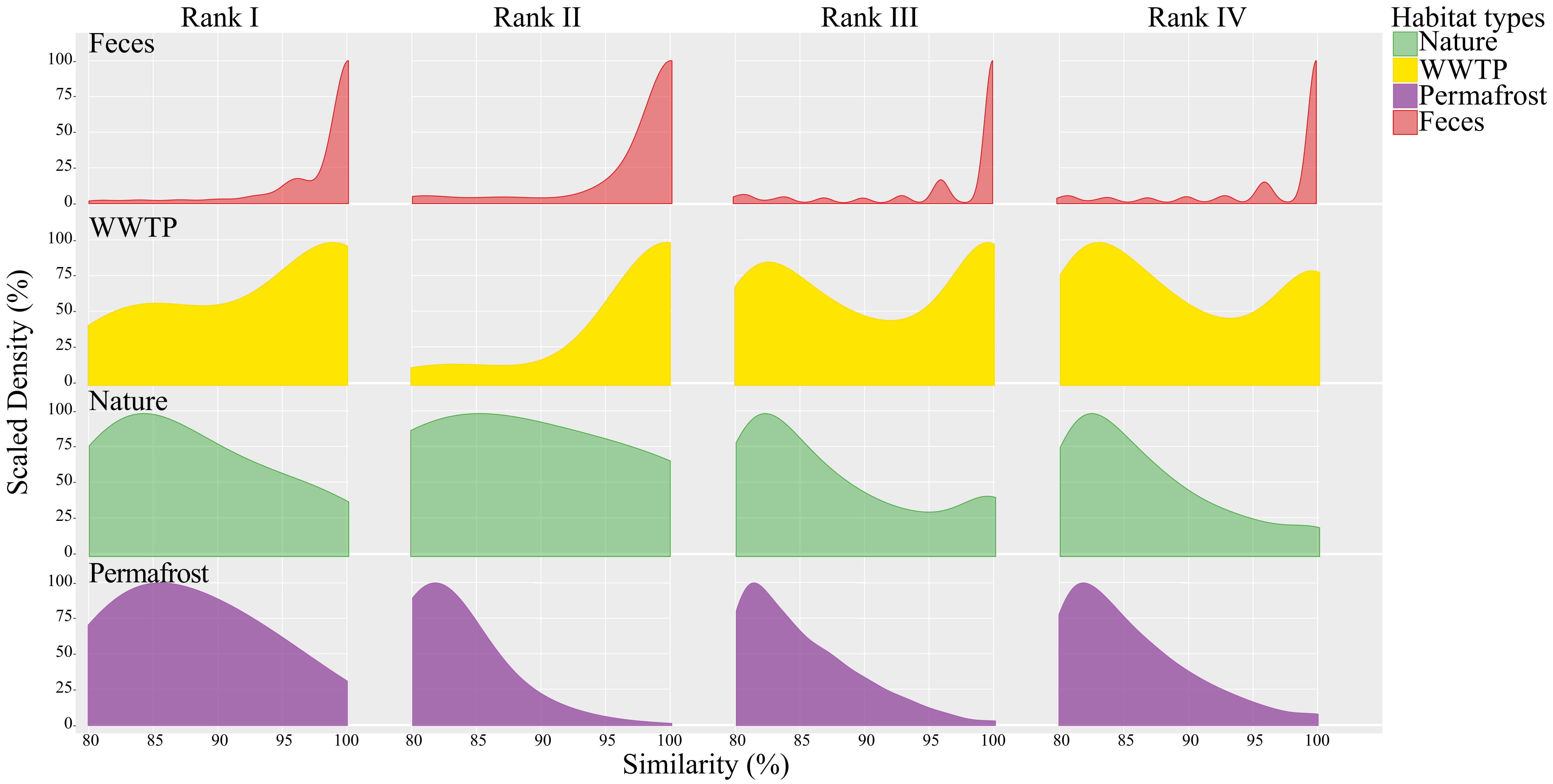

### Supplemental Figure 6

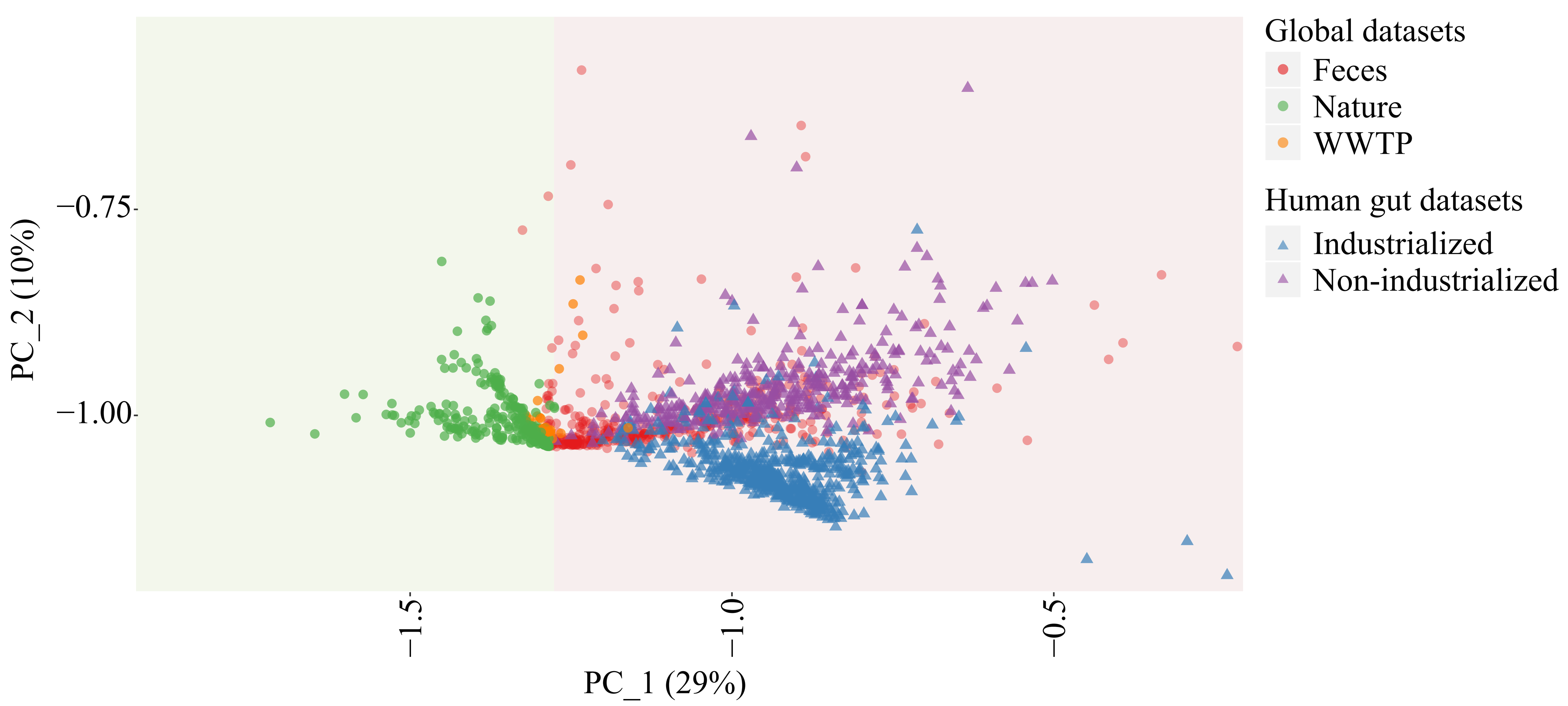

### Supplemental Figure 7

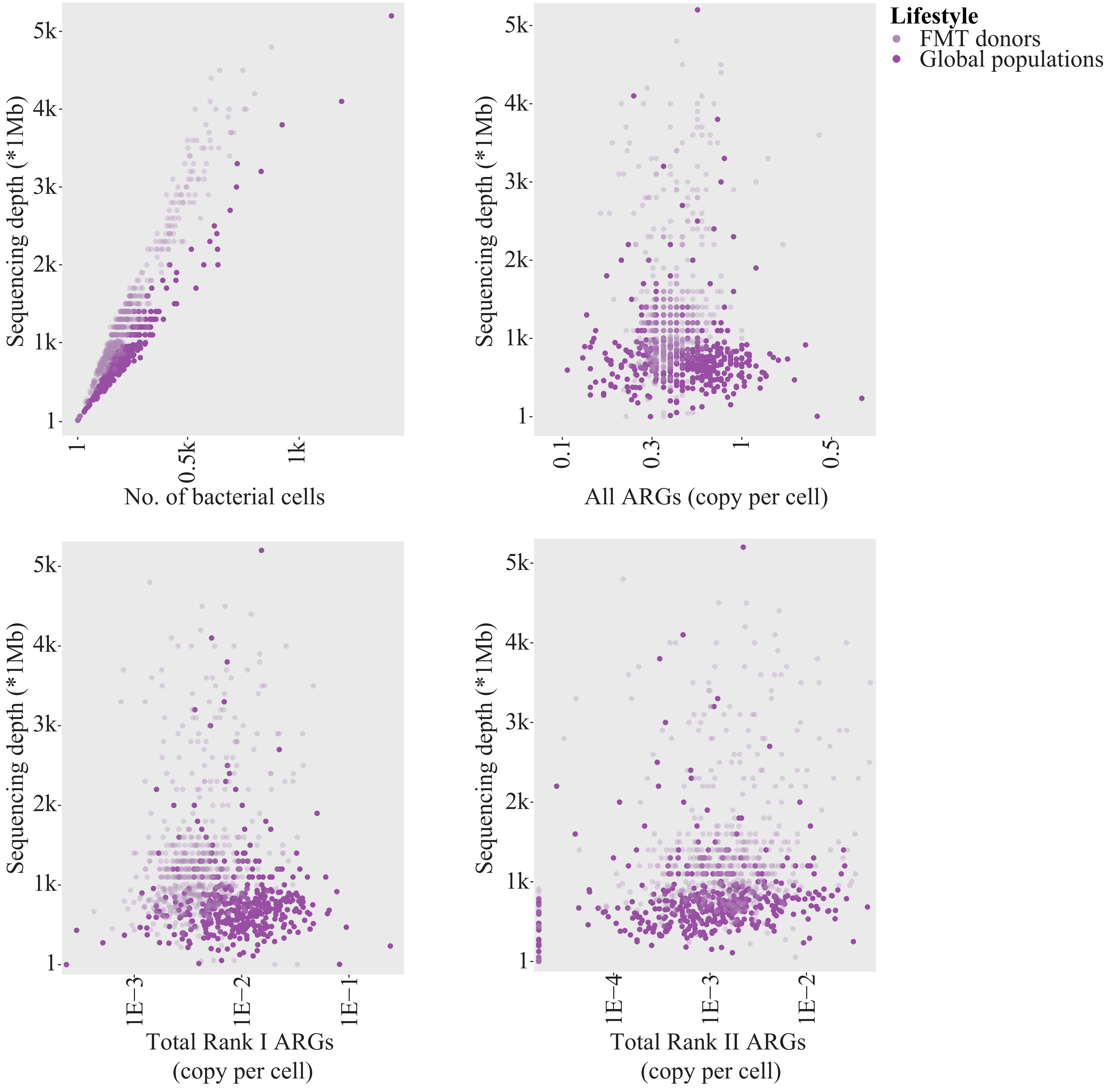

### Supplemental Figure 8

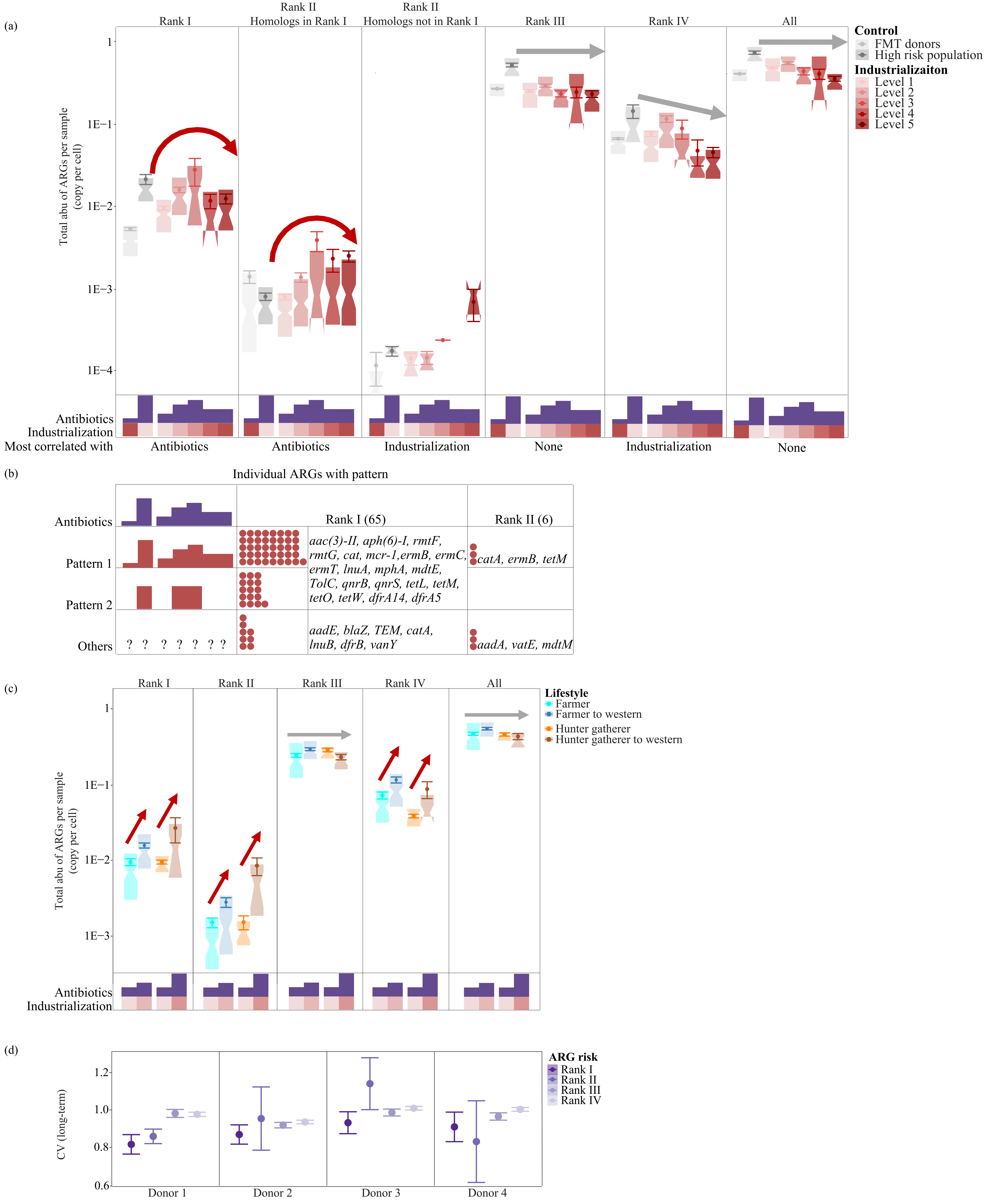
