## Supplemental Table 1 for "Choosing Your Battles: Which Resistance Genes Warrant Global Action?"

| **Table S3.** Total concentration of antibiotics in different habitats collected from literatures reviews. | | | | |
| --- | --- | --- | --- | --- |
| Eco-type | Eco-subtype | Unit | Total Antibiotics | Ref |
| natural_sediment | natural_sediment_China | ng/g | 12.7 | 1 |
| natural_sediment | natural_sediment_China | ng/g | 39.2 | 1 |
| natural_sediment | natural_sediment_China | ng/g | 66.7 | 1 |
| natural_sediment | natural_sediment_gulf | ng/g | 361.4 | 2 |
| natural_sediment | natural_sediment_marine | ng/g | 19.1 | 3 |
| natural_soil | natural_soil_rural | ng/g | 180.1 | 4 |
| natural_soil | natural_soil_rural | ng/g | 51.8 | 5 |
| natural_soil | natural_soil_rural | ng/g | 61.9 | 6 |
| natural_soil | natural_soil_rural | ng/g | 14.0 | 6 |
| natural_soil | natural_soil_rural | ng/g | 69.4 | 7 |
| natural_soil | natural_soil_rural | ng/g | 124.0 | 8 |
| natural_soil | natural_soil_algriculture | ng/g | 923.1 | 9 |
| natural_soil | natural_soil_algriculture | ng/g | 414.7 | 6 |
| natural_soil | natural_soil_algriculture | ng/g | 80.6 | 6 |
| natural_soil | natural_soil_algriculture | ng/g | 30.5 | 6 |
| natural_soil | natural_soil_algriculture | ng/g | 126.9 | 6 |
| natural_soil | natural_soil_algriculture | ng/g | 378.9 | 7 |
| natural_water | natural_water_surfacewater | ng/L | 56.9 | 10 |
| natural_water | natural_water_surfacewater | ng/L | 222.0 | 11 |
| natural_water | natural_water_surfacewater | ng/L | 48.5 | 12 |
| natural_water | natural_water_surfacewater | ng/L | 140.1 | 12 |
| natural_water | natural_water_surfacewater | ng/L | 80.0 | 13 |
| natural_water | natural_water_surfacewater | ng/L | 490.4 | 2 |
| natural_water | natural_water_surfacewater | ng/L | 93.0 | 14 |
| natural_water | natural_water_surfacewater | ng/L | 627.0 | 14 |
| natural_water | natural_water_surfacewater | ng/L | 2560.0 | 14 |
| natural_water | natural_water_surfacewater | ng/L | 209.8 | 15 |
| natural_water | natural_water_surfacewater | ng/L | 61.0 | 16 |
| natural_water | natural_water_surfacewater | ng/L | 113.0 | 16 |
| natural_water | natural_water_surfacewater | ng/L | 133.0 | 16 |
| natural_water | natural_water_surfacewater | ng/L | 147.0 | 16 |
| natural_water | natural_water_surfacewater | ng/L | 360.0 | 16 |
| natural_water | natural_water_surfacewater | ng/L | 77.5 | 17 |
| natural_water | natural_water_surfacewater | ng/L | 648.0 | 18 |
| natural_water | natural_water_surfacewater | ng/L | 2892.4 | 18 |
| natural_water | natural_water_surfacewater | ng/L | 3366.0 | 18 |
| natural_water | natural_water_surfacewater | ng/L | 4135.5 | 18 |
| natural_water | natural_water_surfacewater | ng/L | 6556.0 | 18 |
| natural_water | natural_water_marine | ng/L | 3.4 | 19 |
| natural_water | natural_water_marine | ng/L | 10.4 | 3 |
| natural_water | natural_water_drinking | ng/L | 41.0 | 19 |
| natural_water | natural_water_drinking | ng/L | 346.3 | 19 |
| natural_water | natural_water_coastalwater | ng/L | 1339.0 | 20 |
| wwtp | wwtp_influent | ng/L | 376.5 | 21 |
| wwtp | wwtp_influent | ng/L | 2385.9 | 21 |
| wwtp | wwtp_influent | ng/L | 21900.0 | 22 |
| wwtp | wwtp_influent | ng/L | 3984.0 | 16 |
| wwtp | wwtp_influent | ng/L | 8163.0 | 16 |
| wwtp | wwtp_influent | ng/L | 9649.0 | 16 |
| wwtp | wwtp_influent | ng/L | 160.0 | 16 |
| wwtp | wwtp_effluent | ng/L | 316.2 | 12 |
| wwtp | wwtp_effluent | ng/L | 387.0 | 12 |
| wwtp | wwtp_effluent | ng/L | 648.6 | 12 |
| wwtp | wwtp_effluent | ng/L | 966.1 | 12 |
| wwtp | wwtp_effluent | ng/L | 296.0 | 23 |
| wwtp | wwtp_effluent | ng/L | 879.0 | 23 |
| wwtp | wwtp_effluent | ng/L | 110.0 | 13 |
| wwtp | wwtp_effluent | ng/L | 723.0 | 24 |
| wwtp | wwtp_effluent | ng/L | 201.0 | 16 |
| wwtp | wwtp_effluent | ng/L | 1268.0 | 16 |
| wwtp | wwtp_effluent | ng/L | 8401.0 | 16 |
| wwtp | wwtp_DS | ng/L | 22.6 | 22 |
| wwtp | wwtp_DS | ng/g | 2432.0 | 26 |
| wwtp | wwtp_AS | ng/g | 21508.6 | 27 |
| wwtp | wwtp_effluent | ng/L | 178.1 | 25 |
| fecal_animal | fecal_animal | ng/g | 111217.5 | 9 |
| fecal_animal | fecal_animal | ng/g | 16506.5 | 7 |
| fecal_animal | fecal_animal_chicken | ng/g | 8950.0 | 28 |
| fecal_animal | fecal_animal_chicken | ng/g | 26400.0 | 29 |
| fecal_animal | fecal_animal_pig | ng/g | 27350.0 | 28 |
| fecal_animal | fecal_animal_pig | ng/g | 10121.4 | 8 |
| fecal_animal | fecal_animal_cow | ng/g | 1000.0 | 28 |

**References**

1.	Zhou L-J*, et al.* Trends in the occurrence of human and veterinary antibiotics in the sediments of the Yellow River, Hai River and Liao River in northern China. *Environmental Pollution* **159**, 1877-1885 (2011).

2.	Li W, Shi Y, Gao L, Liu J, Cai Y. Occurrence of antibiotics in water, sediments, aquatic plants, and animals from Baiyangdian Lake in North China. *Chemosphere* **89**, 1307-1315 (2012).

3.	Na G*, et al.* Occurrence, distribution, and bioaccumulation of antibiotics in coastal environment of Dalian, China. *Marine pollution bulletin* **69**, 233-237 (2013).

4.	Xie Y-f*, et al.* Spatial estimation of antibiotic residues in surface soils in a typical intensive vegetable cultivation area in China. *Science of the Total Environment* **430**, 126-131 (2012).

5.	Gibson R, Durán-Álvarez JC, Estrada KL, Chávez A, Cisneros BJ. Accumulation and leaching potential of some pharmaceuticals and potential endocrine disruptors in soils irrigated with wastewater in the Tula Valley, Mexico. *Chemosphere* **81**, 1437-1445 (2010).

6.	Tang X*, et al.* Effects of long-term manure applications on the occurrence of antibiotics and antibiotic resistance genes (ARGs) in paddy soils: evidence from four field experiments in south of China. *Soil Biology and Biochemistry* **90**, 179-187 (2015).

7.	Li C*, et al.* Occurrence of antibiotics in soils and manures from greenhouse vegetable production bases of Beijing, China and an associated risk assessment. *Science of the total environment* **521**, 101-107 (2015).

8.	Aust M-O*, et al.* Distribution of sulfamethazine, chlortetracycline and tylosin in manure and soil of Canadian feedlots after subtherapeutic use in cattle. *Environmental Pollution* **156**, 1243-1251 (2008).

9.	Hu X, Zhou Q, Luo Y. Occurrence and source analysis of typical veterinary antibiotics in manure, soil, vegetables and groundwater from organic vegetable bases, northern China. *Environmental Pollution* **158**, 2992-2998 (2010).

10.	Yoon Y, Ryu J, Oh J, Choi B-G, Snyder SA. Occurrence of endocrine disrupting compounds, pharmaceuticals, and personal care products in the Han River (Seoul, South Korea). *Science of the Total Environment* **408**, 636-643 (2010).

11.	Tamtam F*, et al.* Occurrence and fate of antibiotics in the Seine River in various hydrological conditions. *Science of the Total Environment* **393**, 84-95 (2008).

12.	Zuccato E, Castiglioni S, Bagnati R, Melis M, Fanelli R. Source, occurrence and fate of antibiotics in the Italian aquatic environment. *Journal of hazardous materials* **179**, 1042-1048 (2010).

13.	Bendz D, Paxéus NA, Ginn TR, Loge FJ. Occurrence and fate of pharmaceutically active compounds in the environment, a case study: Höje River in Sweden. *Journal of Hazardous Materials* **122**, 195-204 (2005).

14.	Massey LB, Haggard BE, Galloway JM, Loftin KA, Meyer MT, Green WR. Antibiotic fate and transport in three effluent-dominated Ozark streams. *Ecological Engineering* **36**, 930-938 (2010).

15.	Conley JM, Symes SJ, Schorr MS, Richards SM. Spatial and temporal analysis of pharmaceutical concentrations in the upper Tennessee River basin. *Chemosphere* **73**, 1178-1187 (2008).

16.	Chang X*, et al.* Determination of antibiotics in sewage from hospitals, nursery and slaughter house, wastewater treatment plant and source water in Chongqing region of Three Gorge Reservoir in China. *Environmental pollution* **158**, 1444-1450 (2010).

17.	López-Serna R, Jurado A, Vázquez-Suñé E, Carrera J, Petrović M, Barceló D. Occurrence of 95 pharmaceuticals and transformation products in urban groundwaters underlying the metropolis of Barcelona, Spain. *Environmental Pollution* **174**, 305-315 (2013).

18.	Valcárcel Y, Alonso SG, Rodríguez-Gil J, Gil A, Catalá M. Detection of pharmaceutically active compounds in the rivers and tap water of the Madrid Region (Spain) and potential ecotoxicological risk. *Chemosphere* **84**, 1336-1348 (2011).

19.	Wang Q-J*, et al.* Determination of four fluoroquinolone antibiotics in tap water in Guangzhou and Macao. *Environmental Pollution* **158**, 2350-2358 (2010).

20.	Zou S, Xu W, Zhang R, Tang J, Chen Y, Zhang G. Occurrence and distribution of antibiotics in coastal water of the Bohai Bay, China: impacts of river discharge and aquaculture activities. *Environmental Pollution* **159**, 2913-2920 (2011).

21.	Lin AY-C, Tsai Y-T. Occurrence of pharmaceuticals in Taiwan's surface waters: impact of waste streams from hospitals and pharmaceutical production facilities. *Science of the Total Environment* **407**, 3793-3802 (2009).

22.	Lindberg R, Jarnheimer P-Å, Olsen B, Johansson M, Tysklind M. Determination of antibiotic substances in hospital sewage water using solid phase extraction and liquid chromatography/mass spectrometry and group analogue internal standards. *Chemosphere* **57**, 1479-1488 (2004).

23.	Al Aukidy M, Verlicchi P, Jelic A, Petrovic M, Barcelò D. Monitoring release of pharmaceutical compounds: occurrence and environmental risk assessment of two WWTP effluents and their receiving bodies in the Po Valley, Italy. *Science of the Total Environment* **438**, 15-25 (2012).

24.	Martín J, Camacho-Muñoz D, Santos J, Aparicio I, Alonso E. Occurrence of pharmaceutical compounds in wastewater and sludge from wastewater treatment plants: removal and ecotoxicological impact of wastewater discharges and sludge disposal. *Journal of hazardous materials* **239**, 40-47 (2012).

25.	Ashton D, Hilton M, Thomas K. Investigating the environmental transport of human pharmaceuticals to streams in the United Kingdom. *Science of the Total Environment* **333**, 167-184 (2004).

26.	Lindberg RH, Björklund K, Rendahl P, Johansson MI, Tysklind M, Andersson BA. Environmental risk assessment of antibiotics in the Swedish environment with emphasis on sewage treatment plants. *Water research* **41**, 613-619 (2007).

27.	McClellan K, Halden RU. Pharmaceuticals and personal care products in archived US biosolids from the 2001 EPA national sewage sludge survey. *Water research* **44**, 658-668 (2010).

28.	Hou J*, et al.* Occurrence and distribution of sulfonamides, tetracyclines, quinolones, macrolides, and nitrofurans in livestock manure and amended soils of Northern China. *Environmental Science and Pollution Research* **22**, 4545-4554 (2015).

29.	Dolliver H, Gupta S, Noll S. Antibiotic degradation during manure composting. *Journal of environmental quality* **37**, 1245-1253 (2008).
